# Non-linear gene sets for digital biomarkers of amyotrophic lateral sclerosis

**DOI:** 10.64898/2025.12.18.695103

**Authors:** Keiko Imamura, Ayako Nagahashi, Takuya Yamamoto, Naonori Ueda, Yoshinobu Kawahara, Haruhisa Inoue

## Abstract

Amyotrophic lateral sclerosis (ALS) is a fatal neurodegenerative disease caused by the loss of motor neurons. Accurate and accessible blood-based diagnosis of neurodegenerative diseases, including ALS, is becoming increasingly critical. It would be of a major clinical advantage to be able to distinguish ALS based on gene expression profiles of blood cells, but this still remains to be established. We analyzed existing data of peripheral blood mononuclear cells (PBMCs) of ALS patients by machine learning, the Maximum Mean Discrepancy (MMD), minimizing the issue of multi-collinearity shown in a multiple regression model, and identified a non-linear combination of gene sets to classify healthy controls and ALS, since non-linear approaches can capture intricate gene–gene interactions and threshold effects, which are often critical for accurately distinguishing disease from healthy states. In the gene expression profiles of PBMCs, a combination of expression levels in 3 genes, PRKAR1A, QPCT, and TMEM71, enabled us to classify ALS with an area under the curve (AUC) accuracy of 0.83 from the public database, which were then confirmed by laboratory blood samples with an AUC accuracy of 0.85. Furthermore, we found the expression levels of PRKAR1A, QPCT, and TMEM71 in motor neurons derived from induced pluripotent stem cells (iPSCs), the cell type at the core of the pathology, classified ALS with an accuracy of AUC 0.79. This approach of discriminating ALS with non-linear gene combinations may be useful for identifying ALS molecular biomarkers for blood-based diagnosis as well as for arriving at a completely new perspective of ALS pathogenesis.

## Introduction

Amyotrophic lateral sclerosis (ALS) is a fatal neurodegenerative disease with progressive muscle weakness, atrophy and respiratory failure caused by motor neuron death. Most cases are classified as sporadic ALS, while approximately 10% are familial ALS. ALS diagnosis is based on clinical findings and electrophysiological examinations after the progression of clinical symptoms ^1^. Although the pathological lesions of ALS are located in the brain and spinal cord, biomarkers are being investigated not only in cerebrospinal fluid but also in blood samples. Considerable progress has been made in identifying protein-based liquid biomarkers in the blood, where neurofilaments released from damaged neurons are elevated and reportedly serve as diagnostic, prognostic and monitoring biomarkers of ALS ^2,3^.

Furthermore, there is a growing need for the development of biomarkers that specifically target disease-specific molecular signatures. Peripheral blood mononuclear cells (PBMCs), considered a valuable source for biomarker discovery ^4^, are more accessible than cerebrospinal fluid and, unlike plasma, may enable the detection of low-abundance transcripts and proteins. Whole transcriptome analysis of PBMCs from ALS patients has provided important insights into disease-related gene expression changes ^5^, further supporting their utility in biomarker research. Thus, an approach for discriminating ALS by the data obtained from PBMC samples is promising for a supportive diagnosis of ALS. It would be a major clinical advantage to have access to diagnostics using digital data such as gene expression profiles in blood, which are simple data available from individual cases.

However, due to the complexity of the pathogenesis of ALS, to date, gene expression biomarkers measurable in blood that can support the diagnosis of ALS, especially non-familial, sporadic ALS, have not yet been established. Therefore, it would be useful to identify molecular markers of ALS by conducting analyses via machine learning beyond a human idea-driven approach. Linear models are commonly used in gene expression analyses to identify genes that differ significantly between ALS patients and healthy controls, often linking these genes to disease pathology. Since identifying non-linear combinations of gene expression is crucial because disease mechanisms often involve complex gene–gene interactions that cannot be captured by linear models, we adopted a non-linear analytical framework to better reflect the multifactorial nature of ALS. Using this approach, we aimed to identify a novel set of genes that can discriminate ALS based on blood gene expression profiles. To achieve this, we applied a machine learning method incorporating Maximum Mean Discrepancy (MMD) ^6–8^ to extract gene combinations that distinguish ALS patients from healthy controls.

## Results

To identify combinations of gene sets capable of identifying healthy controls (HC) and amyotrophic lateral sclerosis (ALS) patients, 3D model construction was performed using public gene expression datasets derived from blood samples. Gene sets consisting of ALS causative genes, ALS-related genes, and other genes were analyzed using MMD to extract discriminative gene combinations (Figure 1A). Although a linear model such as linear logistic regression and Hotelling’s *t*^2^ test are generally used for gene expression analysis, we conducted the analysis using a non-linear model because biological phenomena are captured by non-linear science, and the pathogenesis of diseases cannot be explained by a single factor. We introduced MMD ^16,17^, which is capable of detecting non-linear structures in the data-label relationship. The MMD score is shown to be higher if a combination of genes with a difference between healthy control and ALS is selected while, on the other hand, if a combination of genes without a difference between them is selected, the MMD score is shown to be close to 0. In this manner, we extracted genes that classify ALS from healthy control by identifying the combinations that increase the MMD score (Figure 1B).

**Figure 1.**
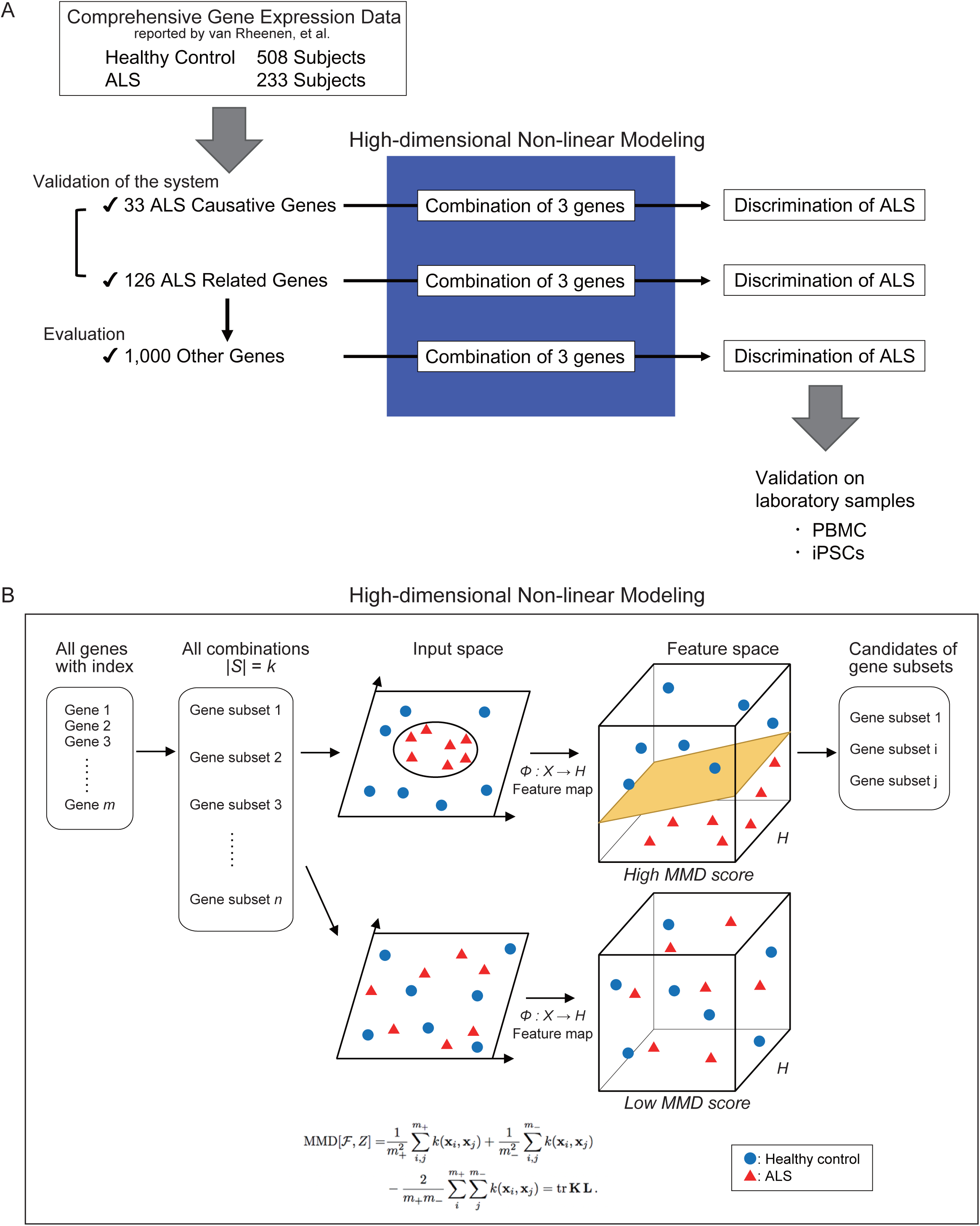
Overview and schematic representation of the analysis pipeline. (A) Schematic diagram of the overall analysis flow in this study. Gene sets consisting of ALS-causative genes, ALS-related genes, and other genes were subjected to analysis using Maximum Mean Discrepancy (MMD) to extract gene combinations that distinguish ALS patients from healthy control. (B) Schematic illustration of the analytical approach using MMD. Non-linear combinatorial analysis, Maximum Mean Discrepancy (MMD), was conducted on gene expression information of blood cells from healthy subjects and ALS subjects to identify gene combinations predicting ALS. The k gene sets were distributed in Hilbert space, and the high accuracy of classifying ALS and healthy control was calculated as MMD score (MMD value) by the non-linear analysis method.

First, as a benchmark, we examined the validity of our approach based on MMD for the extraction of ALS gene sets using a set of genes that is known to be strongly associated with ALS (Figure 2A). Combinations of three genes to classify ALS from healthy controls were selected from genes known to be causative of ALS, which consisted of 33 genes (Figure 2A), and the MMD scores of combinations of three-genes-extracted public datasets were calculated (Figure 2B). We found that the combination of the 3 genes SPG11, CHMP2B, and VCP presented the highest MMD score, which showed an area under the curve (AUC) of 0.75 in receiver operating characteristics (ROC) to classify ALS and healthy subjects (Figure 2C). Next, we investigated the combinations of three genes to classify ALS and healthy control in 126 ALS-related genes (Figure 2A), and their MMD scores were calculated (Figure 2D). The highest MMD score was observed with the combination of CSNK1G3, CHMP2B, and DYNC1H1. The combination of these genes showed an AUC of 0.75 for classifying ALS and healthy control (Figure 2E). These results highlighted the potential of the classification of ALS by MMD using the gene expression data of PBMCs.

**Figure 2.**
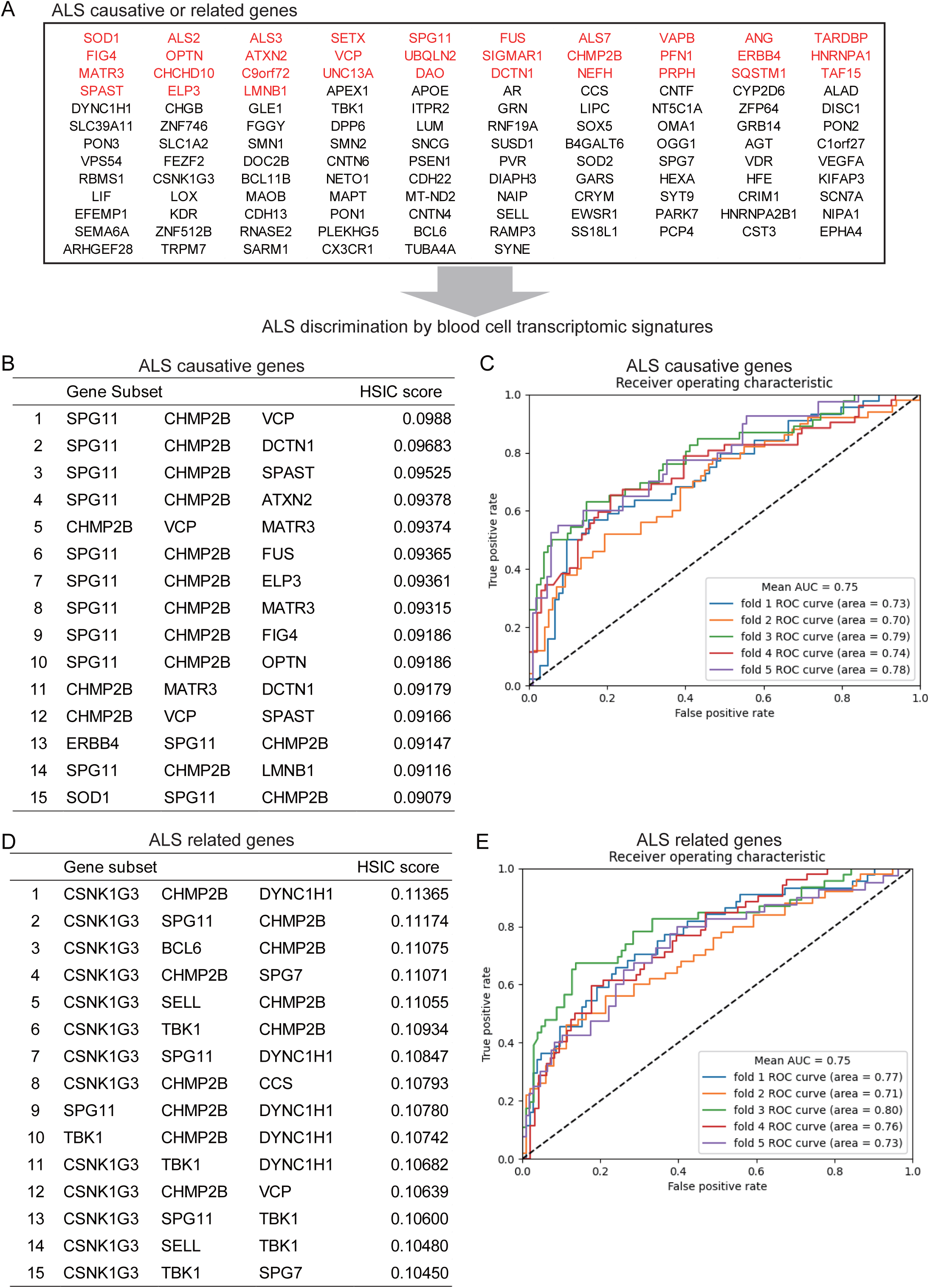
Classification of ALS with ALS-causative/related genes by MMD. A. List of ALS-causative genes and ALS-related genes. Red: ALS-causative genes, Black: ALS-related genes B. Extraction of three genes from ALS-causative genes using MMD. Three genes to classify ALS and healthy control were extracted from the ALS-causative genes using MMD. The top 15 sets are listed. The higher the MMD score the higher the classification accuracy. C: Classification of ALS with 3 ALS-causative genes. ROC to classify ALS and healthy control by combinations of three genes, SPG11, CHMP2B, and VCP, which showed the highest MMD score in Figure 2B. AUC=0.75. D: Extraction of three genes from ALS causative and related genes using MMD. Three genes to classify ALS and healthy control were extracted from ALS-related genes using MMD. The top 15 sets are listed. E: Classification of ALS with 3 ALS-related genes. ROC to classify ALS and healthy control by combinations of three genes, CSNK1G3, CHMP2B, and DYNC1H1, which showed the highest MMD score in Figure 2D. AUC=0.75

In order to investigate unique factors in ALS, we calculated the MMD scores of three-gene combinations from genes not known to be associated with ALS. Figure 3A shows a list of genes sorted by high classification scores of MMD. It is important to avoid the issue of multi-collinearity, an incident with a strong correlation between explanatory variables in a multiple regression model in the selection of genes. Therefore, the combinations of genes that discriminate between healthy control and ALS by logistic regression, a linear model, were also listed (Supplementary Figure 1A), and the most frequent genes were counted; the list of the top 50 combinations included TPT1 (25 appearances), ATP5I (39 appearances), CAPZA2 (17 appearances), and RPL22 (11 appearances) (Supplementary Figure 1B). Among the gene combinations with high MMD scores, we extracted genes that repeatedly appeared, more than 10 times, in the linear regression, and they were excluded. As a result, we found that the combinations of genes with the highest MMD scores were PRKAR1A, QPCT, and TMEM71.

**Figure 3.**
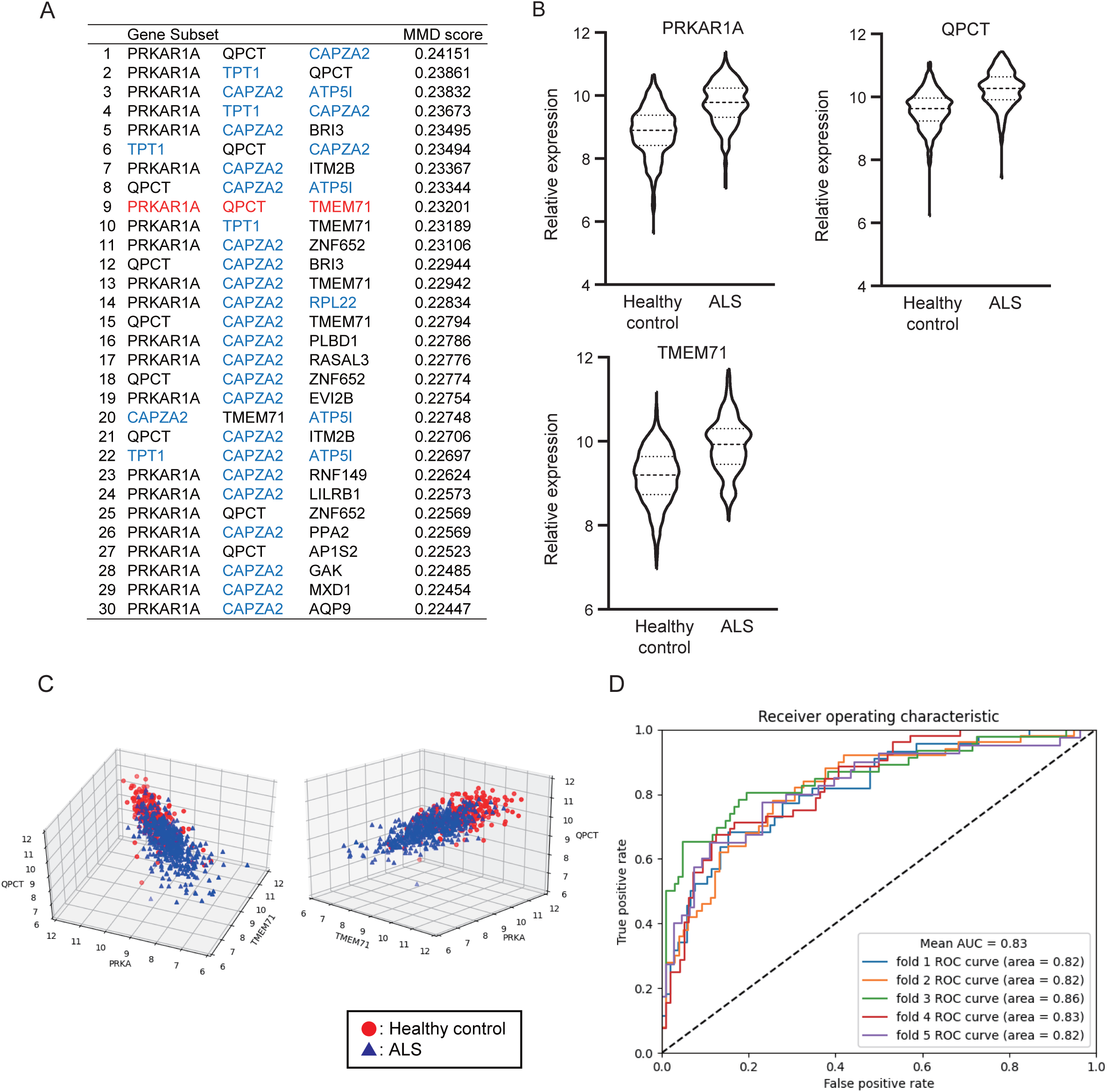
Selection of a gene combination for ALS classification by MMD. A. Three genes were extracted from gene sets except for ALS-causing/related genes to classify ALS and healthy control using MMD. The higher the MMD score the higher the classification accuracy. B. Evaluations of the extracted genes. The expression levels of PRKAR1A, QPCT, and TMEM71 in PBMCs were increased in ALS. n = 508 healthy control; n = 233 ALS. Student-t test, *p < 0.05 C. Three-dimensional analysis using the expression levels of PRKAR1A, QPCT, and TMEM71. Separate distributions of control and ALS are shown. D. ROC for ALS classification by the expression levels of PRKAR1A, QPCT, and TMEM71. AUC=0.83

Next, we verified whether the expression levels of PRKAR1A, QPCT, and TMEM71 enabled us to classify ALS and healthy subjects using public datasets of PBMCs ^9^. The expression levels of each of the three genes were higher in ALS than in controls (Figure 3B). The expression level distributions of the three genes in three dimensions were split between healthy control and ALS (Figure 3C). The algorithm achieved an AUC of 0.83 for the classification of healthy control and ALS (Figure 3D).

In addition, we investigated the association between the expression levels of the three genes and the clinical information on ALS obtained from published data ^9^. The gene expression levels in PRKAR1A and TMEM71 were correlated with survival, although QPCT did not show a significant correlation (Figure 4A). There was no correlation between age of onset and the expression levels of the three genes (Figure 4B). Although there was no difference in QPCT, the expression was significantly higher in the systemic type ALS than in bulbar type ALS, the clinical classification defined by the original data we employed, in PRKAR1A and TMEM71 (Figure 4C).

**Figure 4.**
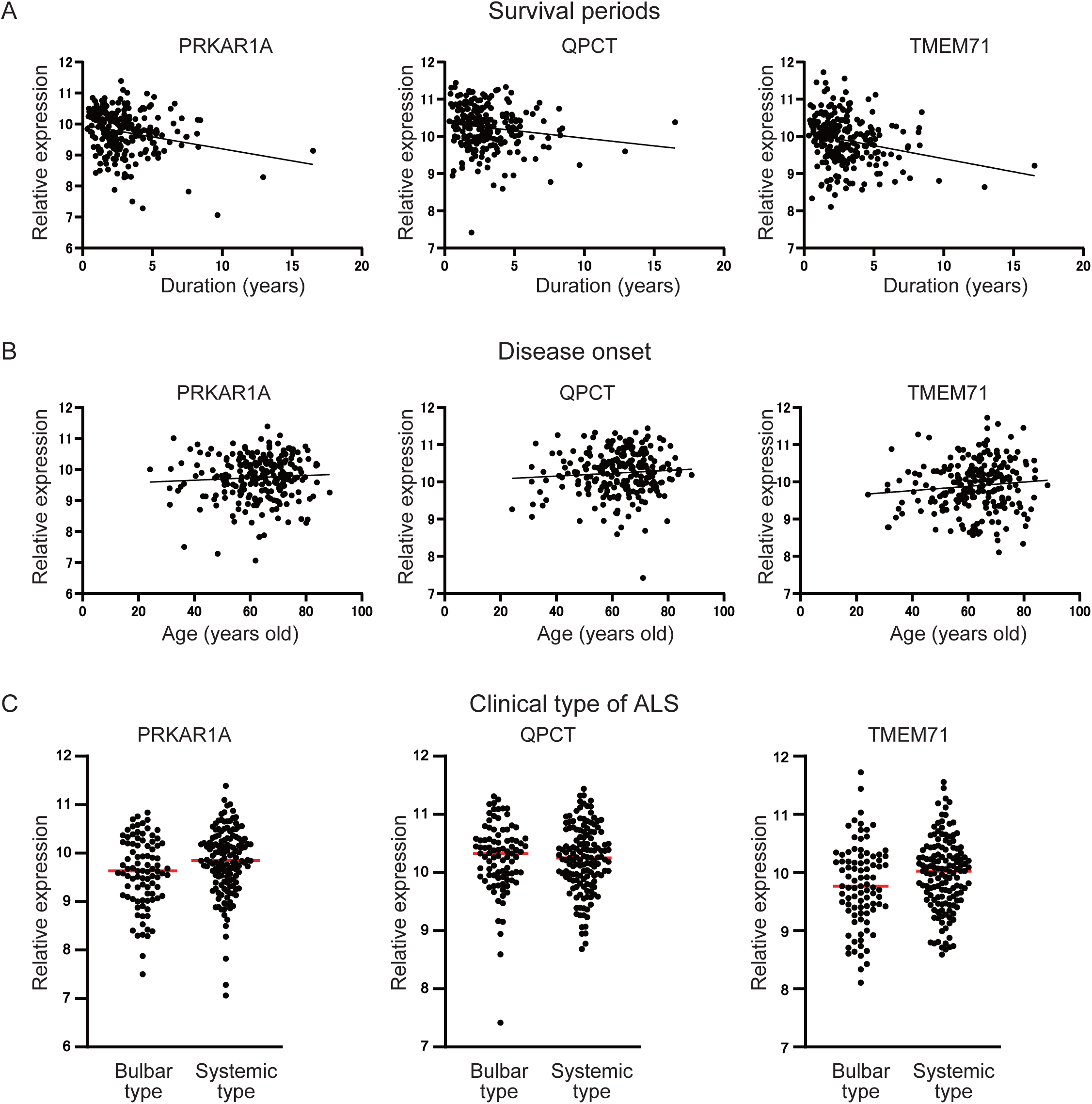
Correlation between gene expression levels and clinical features. A. Correlation of survival period of ALS and gene expression levels. Survival period was correlated with the expression levels of RPKAR1A and TMEM71. B. Correlation of age at onset and gene expression levels. Age at onset of ALS was not correlated with the expression levels of PRKAR1A, QPCT, or TMEM71. C. Relationship between clinical types and gene expression levels. The expression levels of PRKAR1A and TMEM71 were increased in the systemic type of ALS compared to the bulbar type of ALS. p<0.05; n = 142 in systemic type; n = 90 in bulbar type. Student-t test, *p < 0.05

We also examined whether these three genes are classified in other neurodegenerative diseases. Analysis of the blood microarray dataset in Parkinson’s disease ^18^ with 233 healthy controls and 205 idiopathic Parkinson’s disease patients revealed that the AUC of classification of the disease by the three genes was 0.58 (Figure 5A), showing low classification accuracy. Meanwhile, in the dataset of ALS mimics including myelopathy, spinal muscular atrophy, neuropathy etc. ^9^, the AUC for classifying the disease by the three genes was 0.63 (Figure 5B).

**Figure 5.**
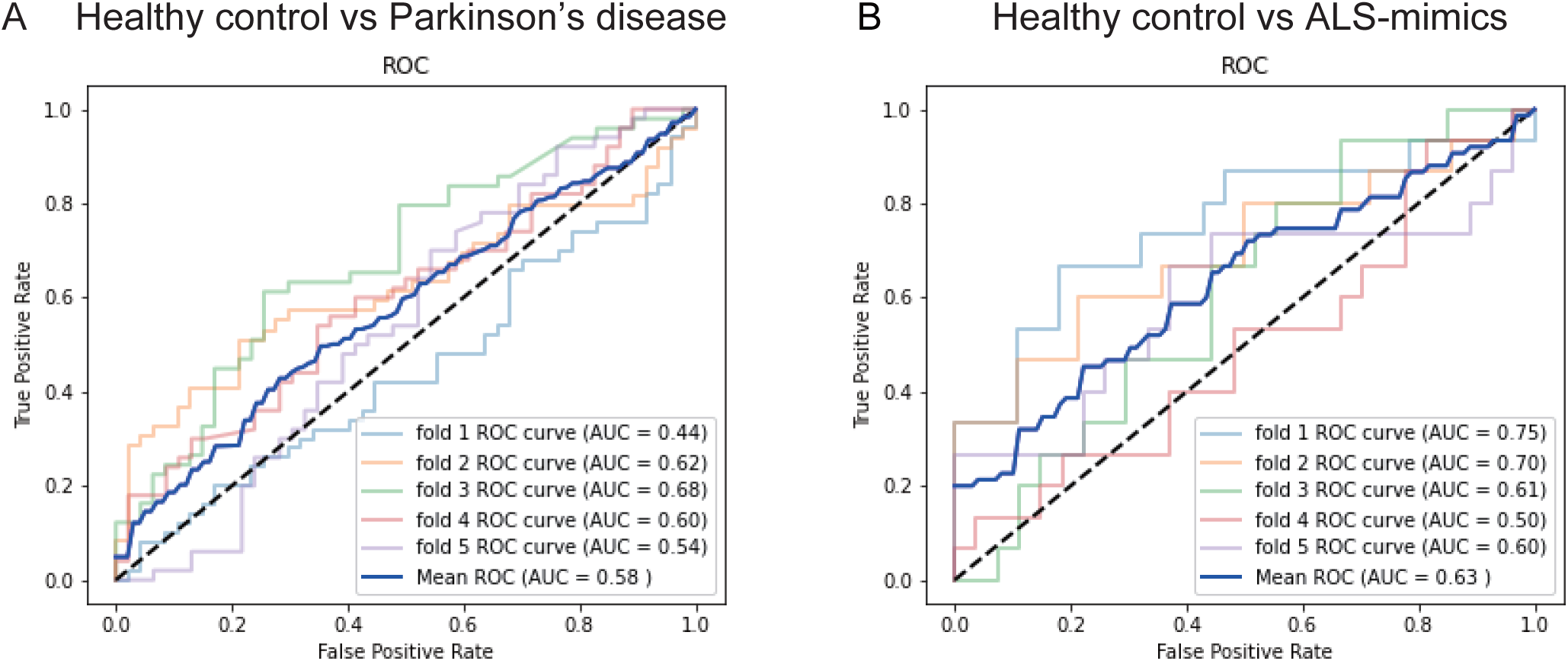
Analysis of other neurological diseases using three-gene combination. A. ROC for classification of Parkinson’s disease by expression levels of PRKAR1A, QPCT, and TMEM71 using the public data GSE99039. AUC=0.58; 233 healthy controls and 205 idiopathic Parkinson’s disease patients. B. ROC for classification of ALS-mimics by the expression levels of PRKAR1A, QPCT, and TMEM71 using public data GSE112680. AUC=0.63; 137 healthy controls and 75 ALS-mimic patients.

To confirm our findings from the database analysis, we investigated the expression levels of the 3 genes, PRKAR1A, QPCT, and TMEM71, using laboratory healthy control PBMCs and ALS patient PBMCs (Figure 6A). PBMCs were collected from 12 ALS patients and 12 healthy subjects, RNA was extracted, and then qPCR analysis was conducted. The expression levels of PRKAR1A and QPCT were significantly higher in ALS compared to healthy control (Figure 6B), while that of TMEM71 showed a tendency to be higher in ALS (Figure 6A). The combination of the expression levels of the three genes led to an AUC of 0.85 for classifying healthy control and ALS (Figure 6C). These results therefore suggested that ALS could be distinguished from healthy control by examining the expression levels of the three genes of PBMCs in real laboratory samples.

**Figure 6.**
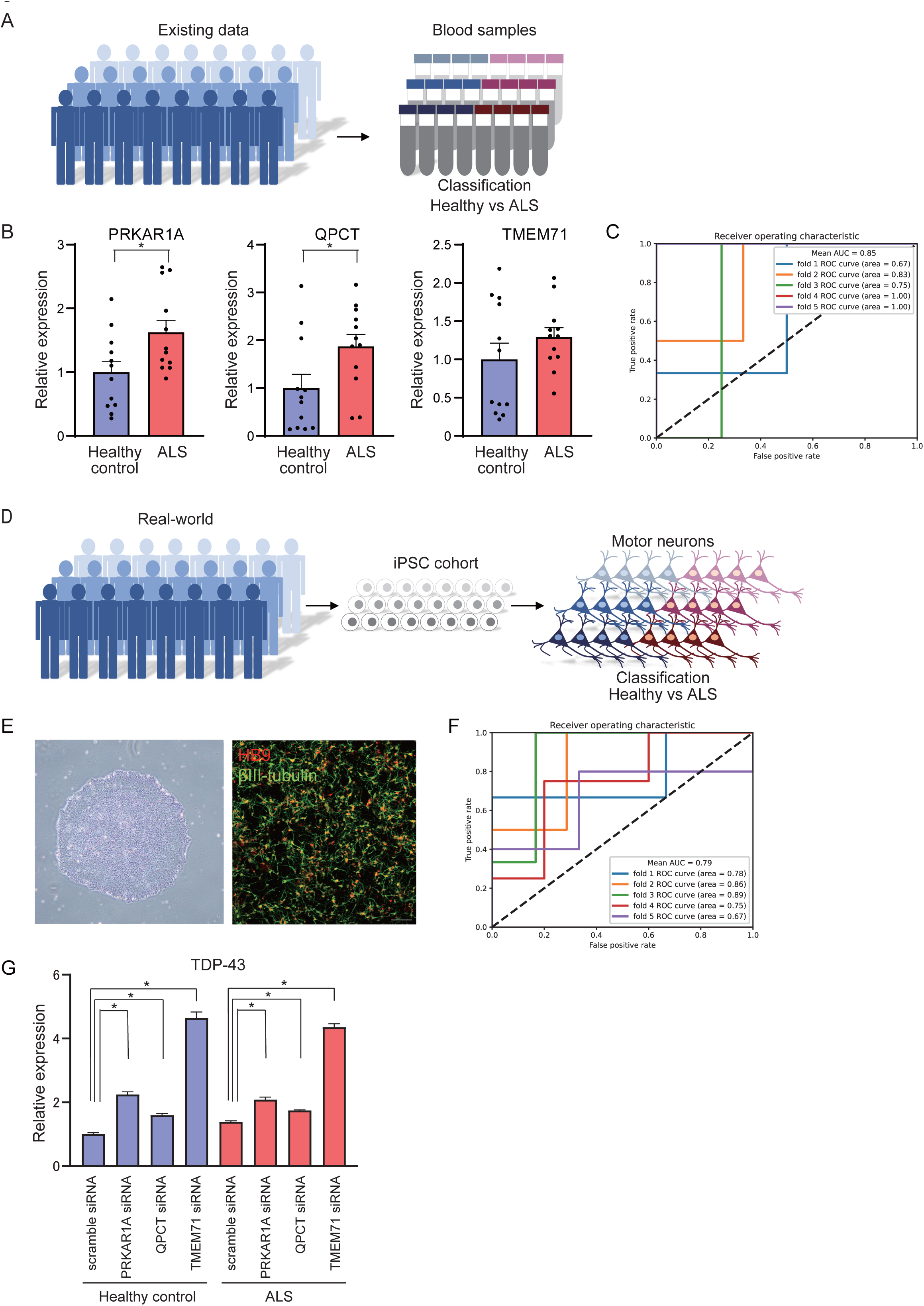
Clinical study for classification of ALS using PBMCs and iPSCs. A. Schema for classifying ALS using PBMCs of laboratory samples. B. Gene expression levels of PRKAR1A, QPCT, and TMEM71 in PBMCs of laboratory samples of healthy subjects and ALS. n = 12 in healthy control; n = 12 in ALS. Student-t test, *p<0.05. C. ROC of ALS classification by expression levels of 3 genes PRKAR1A, QPCT, and TMEM71 using PBMCs. AUC=0.85. D. Schema of clinical study using motor neurons derived from iPSCs. E. iPSCs (left) and motor neurons derived from iPSCs (right). Scale bar = 100 μm. F. Gene expression levels of PRKAR1A, QPCT, and TMEM71 in motor neurons derived from iPSCs. n = 26 in healthy control; n = 18 in ALS. ROC for ALS classification by expression levels of PRKAR1A, QPCT, and TMEM71 using motor neurons derived from iPSCs. AUC=0.79 G. Alteration of TDP-43 mRNA expression by siRNA of PRKAR1A, QPCT, or TMEM71. siRNA of PRKAR1A, QPCT, or TMEM71 increased the expression levels of TDP-43 mRNA in healthy control and ALS motor neurons (n = 3; one-way ANOVA, *p* < 0.005; Dunnett’s *post hoc* test, * *p* < 0.005).

Finally, we examined the expression levels of the three genes in motor neurons derived from iPS cells of 26 healthy controls and 18 ALS patients (Figure 6D, E). Although the expression levels of PRKAR1A, QPCT, or TMEM71 did not differ between motor neurons of healthy control and those of ALS, the combination of the three gene expression levels allowed us to classify healthy control and ALS with an AUC level of 0.79 (Figure 6F). In addition, since TDP-43 accumulation is closely related to the pathogenesis of ALS, the association between the three genes and TDP-43 was investigated. Knock-down of PRKAR1A, QPCT, or TMEM71 by siRNA significantly increased the TDP-43 expression levels in motor neurons in both healthy control and ALS (Figure 6G). These results indicated that the gene set identified by MMD may have some link to the pathogenesis of ALS.

## Discussion

We found blood cell transcriptomic signatures for ALS discrimination using a machine learning algorithm, MMD, a high-dimensional non-linear statistical model. The identified molecular signatures had not previously been focused on ALS. However, we found that siRNA-mediated regulation of the expression of these genes altered the expression levels of TDP-43, a critical, key molecule in ALS, indicating that these molecules could be associated with ALS.

We conducted our analysis using publicly available microarray data derived from blood samples. In a previous study, van Rheenen et al. ^9^, who originally generated this dataset, analyzed gene expression differences between ALS patients and healthy controls. They identified 2,943 differentially expressed transcripts that were primarily involved in RNA binding and intracellular transport. Using these genes, they demonstrated that three machine learning models — support vector machines (SVMs), nearest shrunken centroids (NSC), and LASSO regression — could distinguish ALS patients from healthy individuals with high accuracy. Another study by Swindell et al., ^10^ based on the same dataset, applied SVMs using 850 genes and 468 principal components, and also achieved high classification performance. To identify data not revealed in previous studies, which focused on building classification models, we took another approach by aiming to extract a non-linear set of genes specifically for discriminating ALS since, unlike linear models that consider gene effects independently, non-linear approaches can capture intricate gene–gene interactions and threshold effects, which are often critical for accurately distinguishing disease from healthy states. To maintain independence among the selected genes and avoid multicollinearity, we chose a non-linear model for our analysis.

MMD is used to measure the statistical difference between two random vectors. It then maps the two random vectors in two reproducing kernel Hilbert spaces (RKHS), followed by evaluation of the norm of the means in RKHS to measure their statistical difference between two distributions (23). We applied this model to discover the gene combination to classify ALS and healthy control using blood sample data, as ALS is a heterogeneous disease showing non-linear biological phenomena, and the pathogenesis of the disease cannot be explained by a single factor. By taking advantage of the non-linear model, we succeeded in finding novel combinations of the genes PRKAR1A, QPCT, and TMEM71.

PRKAR1A gene encodes cAMP-dependent protein kinase type I-alpha regulatory subunit, a serine/threonine kinase, which is the main mediator of cAMP signaling in mammals. Phosphorylation mediated by the cAMP/PKA signaling pathway can be elicited by various physiological ligands in cells and is critically involved in the regulation of metabolism, cell proliferation, differentiation, and apoptosis ^19^. Loss of one or both alleles of the gene causes multiple neoplasia syndrome in humans, and loss of both alleles produces an embryonic lethal defect in mice. Although the relationship between the PRKAR1A gene and ALS has still to be clarified, it has been reported that PKA activity is increased in ALS patient spinal cord and SOD1 mice spinal cord ^20^, and that synaptic restoration by cAMP/PKA drives activity-dependent neuroprotection to motoneurons in ALS ^21^. These findings may indicate that increased expression of the PRKAR1A gene might provide a protective effect for ALS.

The QPTC gene encodes Glutaminyl-peptide cyclotransferase. Increased Glutaminyl Cyclase Expression in Peripheral Blood of Alzheimer’s Disease Patients ^22^ and glutaminyl cyclase inhibitors have been reported as a potential Alzheimer’s disease-modifying strategy ^23^. Furthermore, the siRNA screen identifies QPCT as a drug target for Huntington’s disease ^24^, and common polymorphisms of QPCT gene have been associated with the susceptibility for schizophrenia ^25^. Although the association with ALS pathogenesis is as yet unclear, increased expression of the QPCT gene may be associated with a common pathway in neurodegeneration.

TMEM71 encodes a transmembrane protein, although there is yet no clear understanding of its function. Knockout mice present no phenotype and only slight hypothyroidism. TMEM71 expression was increased in glioblastoma and was associated with immune response ^26^, and it also showed a high positive correlation with PD-1 and PD-L1 ^26^. Knockout mice present no phenotype, and only slight hypothyroidism. Increase of the gene expression of TMEM71 may be related to an immune response to ALS.

ALS is a neurodegenerative disease affecting motor neurons in the brain and spinal cord. Although there are several possibilities, including genetic background such as single nucleotide polymorphisms (SNPs) and trafficking between the cerebrospinal and peripheral systems ^27^, the mechanisms responsible for the pathological changes observed in peripheral blood are still unclear. However, the importance of blood biomarkers including the mRNA of blood cells ^9,28^ has been reported in several neurodegenerative diseases ^29,30^, and the development of blood biomarkers is crucial for early diagnosis and timely therapeutic intervention.

We have found a gene combination for discriminating ALS by using the non-linear model MMD and real-world data. Our approach may be useful for identifying PBMC transcriptomic signatures for ALS discrimination as well as for arriving at a completely new perspective of ALS pathogenesis beyond any, until now human idea-driven approach.

## Materials and Methods Study participants

The targeted population with ALS consisted of adults with a diagnosis of “Definite ALS” or “Probable ALS” according to the revised El Escorial criteria. Details of the study participants of healthy subjects and ALS patients for blood collection and iPSC generation are listed in Supplementary table 1 and Supplementary table 2.

### Gene expression data and preprocessing

Gene expression data ^9^ (GSE112676, 233 ALS and 508 healthy control) were used for the MMD analyses. Normalization of gene expression signals was performed according to previously described methods ^10^. Briefly, raw expression intensities and detection p-values (GSE112676_HT12_V3_preQC_nonnormalized.txt) were downloaded and normalized using R limma package (v3.32.10) ^11^ functions (backgroundCorrect and normalizeBetweenArrays). The ComBat algorithm ^12^ implemented in the R sva package (v3.35.2) was used to remove batch effects. The one outlier sample (GSM3077426) even after batch effects correction was excluded from further analyses. ALS-related genes in the ALS online database (ALSoD, https://alsod.ac.uk/) ^13^ were used for MMD prediction in Fig. 1A. Non-biased MMD prediction was performed with the top 1,000 variable genes in detectable expression genes of > 20% of the samples.

### Extraction of gene combinations by Maximum Mean Discrepancy (MMD)

To analyze non-linear structures, we selected MMD ^6^, which has been shown to be significantly advantageous compared to other two-sample test criteria due to its simplicity, interpretability, and the independence from any explicit regularization requirement. For binary classification, a criterion for feature selection can be established by checking whether the distributions are different, and picking a combination of the coordinates of the data that contribute to the difference between the two distributions.

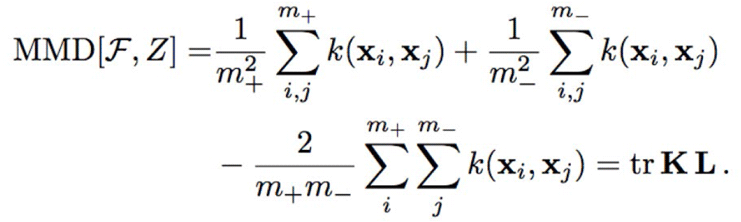

m+ indicates the number of ALS patients, m- indicates the number of healthy control subjects, and x*i* represents the vector of values corresponding to the selected gene for subject *i*. Kernel k employs a Gaussian RBF kernel: k(x, x’) = exp (σ‖||x - x’||^2^). ‖‖ =1/2d, d = |S|: d is the size of the data; selected gene set: S.

Selecting a combination of genes with a difference between ALS and healthy controls results in a higher MMD score, while selecting a combination of genes with no such difference between ALS and healthy controls results in a MMD score close to 0. The combination of genes that maximizes MMD was examined.

### Quantitative RT-PCR

Total RNA of PBMCs or cultured motor neurons was extracted using an RNeasy Plus Mini kit (QIAGEN, location names). One microgram of RNA was reverse-transcribed using ReverTra Ace (TOYOBO, Osaka, Japan). Quantitative PCR analysis was performed using reverse transcription reaction with SYBR Premix Ex TaqⅡ (TAKARA, location names) using StepOnePlus (Thermo Fisher Scientific).

### Generation of iPSCs

iPSCs were generated from fibroblasts or PBMCs of healthy control subjects and ALS patients using episomal vectors for OCT3/4, Sox2, Klf4, L-Myc, Lin28, and dominant-negative p53 or OCT3/4, Sox2, Klf4, L-Myc, Lin28, and shRNA for p53, respectively, as previously reported ^14^. They were cultured by feeder-free and xeno-free culture systems with StemFit (Ajinomoto, location names) with penicillin/streptomycin.

### Motor neuron differentiation from iPSCs

Motor neurons were differentiated from iPSCs as previously described^15^. Briefly, iPSCs were dissociated to single cells and quickly reaggregated in low cell adhesion U-shaped 96-well plates (Lipidule-Coated Plate A-U96, NOF Corporation, Tokyo, Japan). Aggregations were cultured in Dulbecco’s modified Eagle’s medium/Ham’s F12 (Thermo Fisher Scientific, Waltham, MA) containing 5% KSR (Invitrogen, Waltham, MA), minimum essential medium-nonessential amino acids (Invitrogen), L-glutamine (Sigma-Aldrich, St. Louis, MO), 2-mercaptoethanol (Wako, Osaka, Japan), 2 μM dorsomorphin (Sigma-Aldrich), 10 μM SB431542 (Cayman, Ann Arbor, MI), 3 μM CHIR99021 (Cayman), and 12.5 ng/mL fibroblast growth factor (Wako) in a neural inductive stage for 11 days. 100 nM retinoic acid (Sigma-Aldrich) and 500 nM Smoothened ligand (Enzo Life Sciences, Farmingdale, NY) were added on day 4. After patterning with neurobasal medium (Thermo Fisher Scientific) supplemented with B27 Supplement (Thermo Fisher Scientific), 100 nM retinoic acid, 500 nM Smoothened ligand, and 10 μM DAPT (Selleck, Houston, TX), the aggregates were separated by Accumax (Innovative Cell Technologies, San Diego, CA), dissociated into single cells, and adhered to Matrigel (BD Biosciences, Franklin Lakes, NJ)-coated dishes on day 16. Adhesive cells were cultured in neurobasal medium with 10 ng/ml brain-derived neurotrophic factor (R&D Systems, Minneapolis, MN), 10 ng/ml glial cell line-derived neurotrophic factor (R&D Systems), and 10 ng/ml neurotrophin-3 (R&D Systems) for 8 days. On day 21, cells were dissociated to single cells using Accumax and seeded to iMatrix-coated 24-well plates (Corning, location names) at 2×10^5^ cells/well.

### Statistical analysis

Results were analyzed using student’s *t*-test to determine statistical significance. A difference of p < 0.05 was considered significant. Analyses were performed using GraphPad Prism software version 8.0 for Windows (GraphPad Software, San Diego, CA).

## Data availability

The gene expression data of ALS and Parkinson’s disease were obtained from Gene Expression Omnibus (GEO): Accession GSE112681 and GSE99039, respectively. Data supporting the findings of this study are available from the corresponding author upon reasonable request.

## Acknowledgments

We would like to express our sincere gratitude to all of our co-workers and collaborators: Naoya Uematsu, Takako Enami, Kayoko Tsukita, Ichigaku Takigawa for technical support, and Tomomi Urai, Yuki Ueda, Motoko Sugimoto, Chikako Masuda for administrative support.

## Author Contributors

H.I. contributed to the conception and design of the study; K.I, A.N., T.Y, Y.K. contributed to the acquisition and analysis of data; N.U. contributed to supervising; K.I., N.U., Y.K., and H.I. contributed to drafting the manuscript.

## Ethics

The collection and gene expression analysis of PBMCs, and the generation and analysis of human induced pluripotent stem cells (iPSCs) were approved by the Ethics Committees of Kyoto University (the approval number R0091). All methods were performed in accordance with approved guidelines. Formal informed consent was obtained from all subjects. Blood samples were collected from healthy subjects and ALS patients on the basis of their consent.

## References

1. Hardiman, O., van den Berg, L.H., and Kiernan, M.C. (2011). Clinical diagnosis and management of amyotrophic lateral sclerosis. Nat Rev Neurol 7, 639–649. 10.1038/nrneurol.2011.153.

2. Zucchi, E., Bonetto, V., Sorarù, G., Martinelli, I., Parchi, P., Liguori, R., and Mandrioli, J. (2020). Neurofilaments in motor neuron disorders: towards promising diagnostic and prognostic biomarkers. Mol Neurodegener 15, 58. 10.1186/s13024-020-00406-3.

3. Benatar, M., Ostrow, L.W., Lewcock, J.W., Bennett, F., Shefner, J., Bowser, R., Larkin, P., Bruijn, L., and Wuu, J. (2024). Biomarker Qualification for Neurofilament Light Chain in Amyotrophic Lateral Sclerosis: Theory and Practice. Ann Neurol 95, 211–216. 10.1002/ana.26860.

4. Pansarasa, O., Garofalo, M., Scarian, E., Dragoni, F., Garau, J., Di Gerlando, R., Diamanti, L., Bordoni, M., and Gagliardi, S. (2022). Biomarkers in Human Peripheral Blood Mononuclear Cells: The State of the Art in Amyotrophic Lateral Sclerosis. Int J Mol Sci 23. 10.3390/ijms23052580.

5. Dragoni, F., Garofalo, M., Di Gerlando, R., Rizzo, B., Bordoni, M., Scarian, E., Viola, C., Bettoni, V., Fiamingo, G., Tornabene, D., et al. (2025). Whole transcriptome analysis of unmutated sporadic ALS patients’ peripheral blood reveals phenotype-specific gene expression signature. Neurobiol Dis 206, 106823. 10.1016/j.nbd.2025.106823.

6. Borgwardt, K.M., Gretton, A., Rasch, M.J., Kriegel, H.P., Schölkopf, B., and Smola, A.J. (2006). Integrating structured biological data by Kernel Maximum Mean Discrepancy. Bioinformatics 22, e49–57. 10.1093/bioinformatics/btl242.

7. Wang, X., Hu, Z., Yu, T., Wang, Y., Wang, R., Wei, Y., Shu, J., Ma, J., and Li, Y. (2023). Con-AAE: contrastive cycle adversarial autoencoders for single-cell multi-omics alignment and integration. Bioinformatics 39. 10.1093/bioinformatics/btad162.

8. Qian, J., Liao, J., Liu, Z., Chi, Y., Fang, Y., Zheng, Y., Shao, X., Liu, B., Cui, Y., Guo, W., et al. (2023). Reconstruction of the cell pseudo-space from single-cell RNA sequencing data with scSpace. Nat Commun 14, 2484. 10.1038/s41467-023-38121-4.

9. van Rheenen, W., Diekstra, F.P., Harschnitz, O., Westeneng, H.J., van Eijk, K.R., Saris, C.G.J., Groen, E.J.N., van Es, M.A., Blauw, H.M., van Vught, P.W.J., et al. (2018). Whole blood transcriptome analysis in amyotrophic lateral sclerosis: A biomarker study. PLoS One 13, e0198874. 10.1371/journal.pone.0198874.

10. Swindell, W.R., Kruse, C.P.S., List, E.O., Berryman, D.E., and Kopchick, J.J. (2019). ALS blood expression profiling identifies new biomarkers, patient subgroups, and evidence for neutrophilia and hypoxia. J Transl Med 17, 170. 10.1186/s12967-019-1909-0.

11. Ritchie, M.E., Phipson, B., Wu, D., Hu, Y., Law, C.W., Shi, W., and Smyth, G.K. (2015). limma powers differential expression analyses for RNA-sequencing and microarray studies. Nucleic Acids Res 43, e47. 10.1093/nar/gkv007.

12. Leek, J.T., Johnson, W.E., Parker, H.S., Jaffe, A.E., and Storey, J.D. (2012). The sva package for removing batch effects and other unwanted variation in high-throughput experiments. Bioinformatics 28, 882–883. 10.1093/bioinformatics/bts034.

13. Abel, O., Powell, J.F., Andersen, P.M., and Al-Chalabi, A. (2012). ALSoD: A user-friendly online bioinformatics tool for amyotrophic lateral sclerosis genetics. Hum Mutat 33, 1345–1351. 10.1002/humu.22157.

14. Okita, K., Yamakawa, T., Matsumura, Y., Sato, Y., Amano, N., Watanabe, A., Goshima, N., and Yamanaka, S. (2013). An efficient nonviral method to generate integration-free human-induced pluripotent stem cells from cord blood and peripheral blood cells. Stem Cells 31, 458–466. 10.1002/stem.1293.

15. Imamura, K., Yada, Y., Izumi, Y., Morita, M., Kawata, A., Arisato, T., Nagahashi, A., Enami, T., Tsukita, K., Kawakami, H., et al. (2021). Prediction Model of Amyotrophic Lateral Sclerosis by Deep Learning with Patient Induced Pluripotent Stem Cells. Ann Neurol 89, 1226–1233. 10.1002/ana.26047.

16. Zhang, H., Jiang, L., Tang, J., and Ding, Y. (2021). An Accurate Tool for Uncovering Cancer Subtypes by Fast Kernel Learning Method to Integrate Multiple Profile Data. Front Cell Dev Biol 9, 615747. 10.3389/fcell.2021.615747.

17. Liu, Y., Fan, L., Zhang, C., Zhou, T., Xiao, Z., Geng, L., and Shen, D. (2021). Incomplete multi-modal representation learning for Alzheimer’s disease diagnosis. Med Image Anal 69, 101953. 10.1016/j.media.2020.101953.

18. Shamir, R., Klein, C., Amar, D., Vollstedt, E.J., Bonin, M., Usenovic, M., Wong, Y.C., Maver, A., Poths, S., Safer, H., et al. (2017). Analysis of blood-based gene expression in idiopathic Parkinson disease. Neurology 89, 1676–1683. 10.1212/wnl.0000000000004516.

19. Bossis, I., and Stratakis, C.A. (2004). Minireview: PRKAR1A: normal and abnormal functions. Endocrinology 145, 5452–5458. 10.1210/en.2004-0900.

20. Hu, J.H., Chernoff, K., Pelech, S., and Krieger, C. (2003). Protein kinase and protein phosphatase expression in the central nervous system of G93A mSOD over-expressing mice. J Neurochem 85, 422–431. 10.1046/j.1471-4159.2003.01669.x.

21. Bączyk, M., Alami, N.O., Delestrée, N., Martinot, C., Tang, L., Commisso, B., Bayer, D., Doisne, N., Frankel, W., Manuel, M., et al. (2020). Synaptic restoration by cAMP/PKA drives activity-dependent neuroprotection to motoneurons in ALS. J Exp Med 217. 10.1084/jem.20191734.

22. Valenti, M.T., Bolognin, S., Zanatta, C., Donatelli, L., Innamorati, G., Pampanin, M., Zanusso, G., Zatta, P., and Dalle Carbonare, L. (2013). Increased glutaminyl cyclase expression in peripheral blood of Alzheimer’s disease patients. J Alzheimers Dis 34, 263–271. 10.3233/jad-120517.

23. Vijayan, D., and Chandra, R. (2020). Amyloid Beta Hypothesis in Alzheimer’s Disease: Major Culprits and Recent Therapeutic Strategies. Curr Drug Targets 21, 148–166. 10.2174/1389450120666190806153206.

24. Jimenez-Sanchez, M., Lam, W., Hannus, M., Sönnichsen, B., Imarisio, S., Fleming, A., Tarditi, A., Menzies, F., Dami, T.E., Xu, C., et al. (2015). siRNA screen identifies QPCT as a druggable target for Huntington’s disease. Nat Chem Biol 11, 347–354. 10.1038/nchembio.1790.

25. Zhang, Q.Q., Jiang, T., Gu, L.Z., Zhu, X.C., Zhao, H.D., Gao, Q., Zhu, H.Q., Zhou, J.S., and Zhang, Y.D. (2016). Common Polymorphisms Within QPCT Gene Are Associated with the Susceptibility of Schizophrenia in a Han Chinese Population. Mol Neurobiol 53, 6362–6366. 10.1007/s12035-015-9541-3.

26. Wang, K.Y., Huang, R.Y., Tong, X.Z., Zhang, K.N., Liu, Y.W., Zeng, F., Hu, H.M., and Jiang, T. (2019). Molecular and clinical characterization of TMEM71 expression at the transcriptional level in glioma. CNS Neurosci Ther 25, 965–975. 10.1111/cns.13137.

27. Komiya, H., Takeuchi, H., Ogawa, Y., Hatooka, Y., Takahashi, K., Katsumoto, A., Kubota, S., Nakamura, H., Kunii, M., Tada, M., et al. (2020). CCR2 is localized in microglia and neurons, as well as infiltrating monocytes, in the lumbar spinal cord of ALS mice. Mol Brain 13, 64. 10.1186/s13041-020-00607-3.

28. Augustine, J., and Jereesh, A.S. (2022). Blood-based gene-expression biomarkers identification for the non-invasive diagnosis of Parkinson’s disease using two-layer hybrid feature selection. Gene 823, 146366. 10.1016/j.gene.2022.146366.

29. Shefner, J.M., Bedlack, R., Andrews, J.A., Berry, J.D., Bowser, R., Brown, R., Glass, J.D., Maragakis, N.J., Miller, T.M., Rothstein, J.D., and Cudkowicz, M.E. (2022). Amyotrophic Lateral Sclerosis Clinical Trials and Interpretation of Functional End Points and Fluid Biomarkers: A Review. JAMA Neurol 79, 1312–1318. 10.1001/jamaneurol.2022.3282.

30. Teunissen, C.E., Verberk, I.M.W., Thijssen, E.H., Vermunt, L., Hansson, O., Zetterberg, H., van der Flier, W.M., Mielke, M.M., and Del Campo, M. (2022). Blood-based biomarkers for Alzheimer’s disease: towards clinical implementation. Lancet Neurol 21, 66–77. 10.1016/s1474-4422(21)00361-6.

