## Supplementary file for "Non-linear gene sets for digital biomarkers of amyotrophic lateral sclerosis"

**Title: ALS-Discriminative Gene Combinations in Blood Cells Revealed by Computational Analysis**

**Supplementary Figure 1. Analysis of gene sets from linear regression model**

**Supplementary table 1. List of PBMCs**

**Supplementary table 2. List of iPSCs**

Supplementary Figure 1

A

| | Gene Subset | | | AUC |
|---|---|---|---|---|
| 1 | TPT1 | TMEM71 | ATP5I | 0.895767 |
| 2 | TPT1 | ATP5I | VNN2 | 0.889306 |
| 3 | TPT1 | ATP5I | BRI3 | 0.888054 |
| 4 | TPT1 | RPL22 | VNN2 | 0.884817 |
| 5 | TPT1 | TMEM71 | RPL22 | 0.884791 |
| 6 | TPT1 | RPL22 | SARAF | 0.883219 |
| 7 | TPT1 | ATP5I | RNF149 | 0.882393 |
| 8 | TPT1 | RNF149 | RPL22 | 0.881793 |
| 9 | TPT1 | ATP5I | PLBD1 | 0.881407 |
| 10 | TPT1 | ATP5I | AQP9 | 0.881367 |
| 11 | TPT1 | ATP5I | MXD1 | 0.881141 |
| 12 | TPT1 | CAPZA2 | ATP5I | 0.880941 |
| 13 | CAPZA2 | TMEM71 | ATP5I | 0.880368 |
| 14 | TMEM71 | RPL22 | GAK | 0.880288 |
| 15 | CAPZA2 | ATP5I | IL2RB | 0.879862 |
| 16 | TPT1 | QPCT | ATP5I | 0.879249 |
| 17 | CAPZA2 | ATP5I | LILRB1 | 0.879156 |
| 18 | TPT1 | RASAL3 | RPL22 | 0.879049 |
| 19 | TPT1 | BRI3 | RPL22 | 0.878796 |
| 20 | RPL22 | SARAF | IL2RB | 0.878663 |
| 21 | CAPZA2 | ATP5I | MATK | 0.878117 |
| 22 | PRKAR1A | TPT1 | ATP5I | 0.877904 |
| 23 | CAPZA2 | ATP5I | RASAL3 | 0.877771 |
| 24 | TPT1 | ATP5I | RASAL3 | 0.877145 |
| 25 | CAPZA2 | ATP5I | GAK | 0.876985 |
| 26 | CAPZA2 | ATP5I | BRI3 | 0.876958 |
| 27 | CAPZA2 | ATP5I | EIF3B | 0.876758 |
| 28 | CAPZA2 | ATP5I | VNN2 | 0.876758 |
| 29 | CAPZA2 | ATP5I | ZNF652 | 0.876638 |
| 30 | CAPZA2 | ATP5I | PLBD1 | 0.876252 |

B

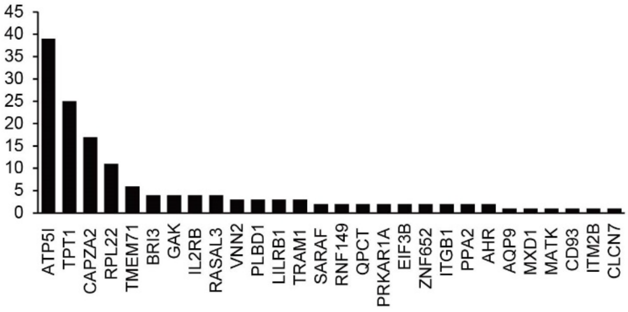


C

| | Gene Subset | | | | MMD score |
|---|---|---|---|---|---|
| 1 | PRKAR1A | QPCT | TMEM71 | CAPZA2 | 0.23516 |
| 2 | PRKAR1A | QPCT | TMEM71 | TPT1 | 0.23255 |
| 3 | PRKAR1A | QPCT | TMEM71 | - | 0.23201 |
| 4 | PRKAR1A | QPCT | TMEM71 | ZNF652 | 0.22405 |
| 5 | PRKAR1A | QPCT | TMEM71 | ATP5I | 0.22290 |
| 6 | PRKAR1A | QPCT | TMEM71 | BRI3 | 0.22264 |
| 7 | PRKAR1A | QPCT | TMEM71 | ITM2B | 0.22192 |
| 8 | PRKAR1A | QPCT | TMEM71 | TMEM123 | 0.22110 |
| 9 | PRKAR1A | QPCT | TMEM71 | AP1S2 | 0.22041 |
| 10 | PRKAR1A | QPCT | TMEM71 | PTBP3 | 0.21963 |

**Supplementary Figure 1. Analysis of gene sets from linear regression model**

A. Extraction of genes to distinguish ALS from healthy control using linear regression.

B. Frequencies of genes appearing repeatedly in the top 50 of the linear model.

C. Diagnostic classification of ALS based on a combination of four genes.

**Supplementary table 1. List of PBMC**

| | PBMC clone | Gender | Duration of illness | ALS-related mutation | ALS type |
|---|---|---|---|---|---|
| Healthy control | hc10 | M | N.A. | N.A. | N.A. |
| | hc12 | M | N.A. | N.A. | N.A. |
| | hc18 | F | N.A. | N.A. | N.A. |
| | hc19 | M | N.A. | N.A. | N.A. |
| | hc20 | M | N.A. | N.A. | N.A. |
| | hc29 | F | N.A. | N.A. | N.A. |
| | hc32 | F | N.A. | N.A. | N.A. |
| | hc37 | M | N.A. | N.A. | N.A. |
| | hc40 | M | N.A. | N.A. | N.A. |
| | hc46 | F | N.A. | N.A. | N.A. |
| | hc47 | F | N.A. | N.A. | N.A. |
| | hc50 | F | N.A. | N.A. | N.A. |
| Sporadic ALS | ALS74 | M | 0.5 | none | systemic |
| | ALS79 | M | 3.7 | none | systemic |
| | ALS81 | F | 18 | none | systemic |
| | ALS89 | F | 7.0 | none | systemic |
| | ALS91 | F | 3.0 | none | systemic |
| | ALS128 | M | 0.8 | none | systemic |
| | ALS137 | F | 0.7 | none | bulbar |
| | ALS149 | M | 1.3 | none | systemic |
| | ALS150 | M | 1.5 | none | systemic |
| | ALS151 | M | 2.5 | none | systemic |
| | ALS153 | M | 1.6 | none | systemic |
| | ALS154 | M | 3.3 | none | systemic |

**Supplementary table 2. List of iPSCs**

| | iPSC clone | Gender | Duration of illness | ALS-related mutation | ALS type | Origin | Reprogramming |
|---|---|---|---|---|---|---|---|
| Healthy control | 201B7 | F | N.A. | N.A. | N.A. | fibroblast | retrovirus |
| | N112E14 | F | N.A. | N.A. | N.A. | PBMC | episomal |
| | hc1F | M | N.A. | N.A. | N.A. | PBMC | episomal |
| | hc3E | F | N.A. | N.A. | N.A. | PBMC | episomal |
| | hc2EL1 | M | N.A. | N.A. | N.A. | PBMC | episomal |
| | hc4EL2 | F | N.A. | N.A. | N.A. | PBMC | episomal |
| | hc12EL1 | M | N.A. | N.A. | N.A. | PBMC | episomal |
| | hc13EL1 | F | N.A. | N.A. | N.A. | PBMC | episomal |
| | hc17EL3 | M | N.A. | N.A. | N.A. | PBMC | episomal |
| | hc18EL7 | F | N.A. | N.A. | N.A. | PBMC | episomal |
| | hc22EL1 | F | N.A. | N.A. | N.A. | PBMC | episomal |
| | hc23EL1 | M | N.A. | N.A. | N.A. | PBMC | episomal |
| | hc5EL2 | F | N.A. | N.A. | N.A. | PBMC | episomal |
| | hc6B | M | N.A. | N.A. | N.A. | PBMC | episomal |
| | hc9EL4 | F | N.A. | N.A. | N.A. | PBMC | episomal |
| | hc10EL1 | M | N.A. | N.A. | N.A. | PBMC | episomal |
| | hc14EL10 | M | N.A. | N.A. | N.A. | PBMC | episomal |
| | hc15EL7 | F | N.A. | N.A. | N.A. | PBMC | episomal |
| | hc16EL4 | F | N.A. | N.A. | N.A. | PBMC | episomal |
| | hc19EL1 | M | N.A. | N.A. | N.A. | PBMC | episomal |
| | hc20EL1 | M | N.A. | N.A. | N.A. | PBMC | episomal |
| | hc24EL1 | F | N.A. | N.A. | N.A. | PBMC | episomal |
| | hc11EL5 | F | N.A. | N.A. | N.A. | PBMC | episomal |
| | hc8EL4 | M | N.A. | N.A. | N.A. | PBMC | episomal |
| | hc25EL3 | M | N.A. | N.A. | N.A. | PBMC | episomal |
| | hc26EL1 | F | N.A. | N.A. | N.A. | PBMC | episomal |
| Sporadic ALS | A13-1 | F | 2 | none | systemic | fibroblast | episomal |
| | ALS1F | M | 3 | none | systemic | fibroblast | episomal |
| | ALS3/7C | M | 1 | none | systemic | fibroblast | episomal |
| | ALS12F | M | 2 | none | systemic | fibroblast | episomal |
| | ALS17E | F | 1 | none | bulbar | fibroblast | episomal |
| | ALS19C | F | 1 | none | systemic | fibroblast | episomal |
| | ALS23F | F | 3 | none | bulbar | fibroblast | episomal |

| | | | | | | | |
|---|---|---|---|---|---|---|---|
| | ALS38F | M | 1 | none | systemic | fibroblast | episomal |
| | ALS66E | F | 4 | none | systemic | PBMC | episomal |
| | ALS71F | M | 4 | none | systemic | PBMC | episomal |
| | ALS72F | M | 8 | none | systemic | PBMC | episomal |
| | ALS74F | M | 0.5 | none | systemic | PBMC | episomal |
| | ALS85E | F | 8 | none | bulbar | PBMC | episomal |
| | ALS86E | M | 13 | none | systemic | PBMC | episomal |
| | ALS88E | F | 17 | none | systemic | PBMC | episomal |
| | ALS89F | F | 7 | none | systemic | PBMC | episomal |
| | ALS90E | F | 2 | none | systemic | PBMC | episomal |
| | ALS91E | F | 3 | none | systemic | PBMC | episomal |
